# Determining potential immunomodulatory drug efficacy in sepsis using ELISpot

**DOI:** 10.1101/2024.07.10.602970

**Authors:** AH Walton, MB Mazer, KE Remy, EB Davitt, TS Griffith, RW Gould, VP Badovinac, SC Brakenridge, AM Drewry, TJ Loftus, PA Efron, LL Moldawer, CC Caldwell, RS Hotchkiss

## Abstract

**Purpose:** This study evaluated the ability of ELISpot to identify immuno-modulatory drug therapies for their potential efficacy in patients with sepsis.

**Methods:** ELISpot was performed using diluted whole blood from 61 septic patients and 48 healthy matched controls. Innate and adaptive immunity were evaluated by *ex vivo* stimulated production of TNF-α and IFN-γ respectively. Potential drug efficacy was determined by the drugs’ effects to increase or decrease the number of cytokine-producing cells and amount of cytokine produced per cell as determined by spot size and intensity. The corticosteroid dexamethasone was evaluated for its ability to down modulate TNF-α and IFN-γ production. The TLR7/8 agonist resiquimod (R848) and T-cell stimulants IL-7 and anti-PD-1 mAb were tested for their ability to enhance immune responses in sepsis.

**Results:** Spontaneous production of TNF-α and IFN-γ varied among healthy subjects and septic patients. LPS or resiquimod stimulation increased total TNF-α production in septic patients by 1,648% and 1,929% respectively. Conversely, dexamethasone diminished the responses to LPS or resiquimod by 71% and 61% respectively. IL-7, but not anti-PD-1 mAb markedly increased IFN-γ production in both healthy subjects (127%) and septic patients (79%). Dexamethasone also reduced anti-CD3/CD28 mAb stimulated IFN-γ production by 54%; while IL-7 ameliorated dexamethasone-induced suppression. IL-7 significantly enhanced lymphocyte function in over 90% of septic patients.

**Conclusion:** ELISpot can reveal host immune response patterns and the effects of drugs to selectively down– or up-regulate patient immunity. Furthermore, the ability of ELISpot to detect the effect of specific immuno-modulatory drugs to independently regulate the innate and adaptive host response could enable precision-based immune drug therapies in sepsis.

## INTRODUCTION

Sepsis is life-threatening organ dysfunction caused by a dysregulated host response to infection (1). An abnormal immune response to the invading pathogens is undoubtedly a major pathophysiologic abnormality driving the dysregulated host response (2–4). Sepsis frequently progresses from an initial potentially damaging hyper-inflammatory response to a more prolonged and sustained immunosuppressive phase which may compromise the ability of the patient to eradicate primary pathogens and/or render the patient more susceptible to secondary infections (3,4). Previous clinical trials have tested a variety of immune modulatory drugs that can either antagonize specific inflammatory molecules, suppress systemic inflammation or conversely, reverse sepsis-induced immune suppression (5–9). Currently, no sepsis drug therapies have consistently improved patient outcomes. A key stumbling block when testing immune modulatory drug therapies in sepsis has been an inability to more precisely target the correct intervention to the functional state of the individual septic patient’s immune system at the time of intervention (8–10).

Recent work from our group found sepsis mortality could be accurately predicted using the results of *ex vivo* T cell receptor stimulation of whole blood production of interferon-gamma (IFN-γ) using the enzyme-linked immunospot (ELISpot) assay, as the septic patients who died had an impaired adaptive host immune response as early as day 1-4 of hospital admission (11). Both the number of IFN-γ producing cells and the amount of IFN-γ produced per lymphocyte was decreased in patients who ultimately succumbed of sepsis. In more recent studies, sepsis patients who had *both* reduced *ex vivo* IFN-γ and tumor necrosis factor-alpha (TNF-α) production by ELISpot had markedly increased risk of death compared to sepsis patients whose responses were greater than the median response by healthy subjects (*manuscript submitted*). These findings are potentially very important and, if confirmed in subsequent larger studies, suggest that ELISpot can be useful in identifying those septic patients who are more profoundly immune suppressed and have an increased risk for mortality, thereby making them ideal candidates for adjuvant drug therapies that restore host immunity. In addition to identifying septic patients who are appropriate candidates for therapies to up-regulate host immunity, ELISpot may also be useful in determining whether patients with sepsis are manifesting an excessive hyper-inflammatory response to the septic insult and would benefit from drugs that *down modulate* the immune response.

The purpose of the present study was to determine to what extent a diluted whole blood ELISpot assay could be useful in guiding potential immune modulatory therapies in patients with sepsis by identifying which drugs are most active in augmenting host immunity in *ex vivo* blood samples from patients with sepsis. We tested drugs that act to either suppress a putative hyper-immune response or to reverse sepsis-induced immunosuppression. To increase the clinical relevance of our studies, with the exception of LPS, we employed drugs that are either FDA approved or in active clinical trials. Finally, we tested drugs that act on both innate and adaptive immune effector cells to gain potential mechanistic insight into sepsis-induced defects in patients.

## METHODS

### Patient enrollment

Patient enrollment was carried out at seven sites across the US as a part of the SPIES consortium as previously described (11). Inclusion criteria consisted primarily of ICU admission with sepsis from either severe trauma, non-trauma, postoperative ICU admission, ICU transfer from the emergency department, or inpatient transfer from ward to ICU. A small proportion of patients were admitted directly from the emergency department with suspicion of community-acquired sepsis. Sepsis was defined based on the Sepsis-3 criteria (1) and all participants were clinically adjudicated at the individual participating sites.

Healthy control participants were recruited at each of the clinical sites. Efforts were made to match the healthy control participants’ age, sex, and race/ethnicity to those of the sepsis cohort. (**Tables 1 & 2**).

**Table 1:**
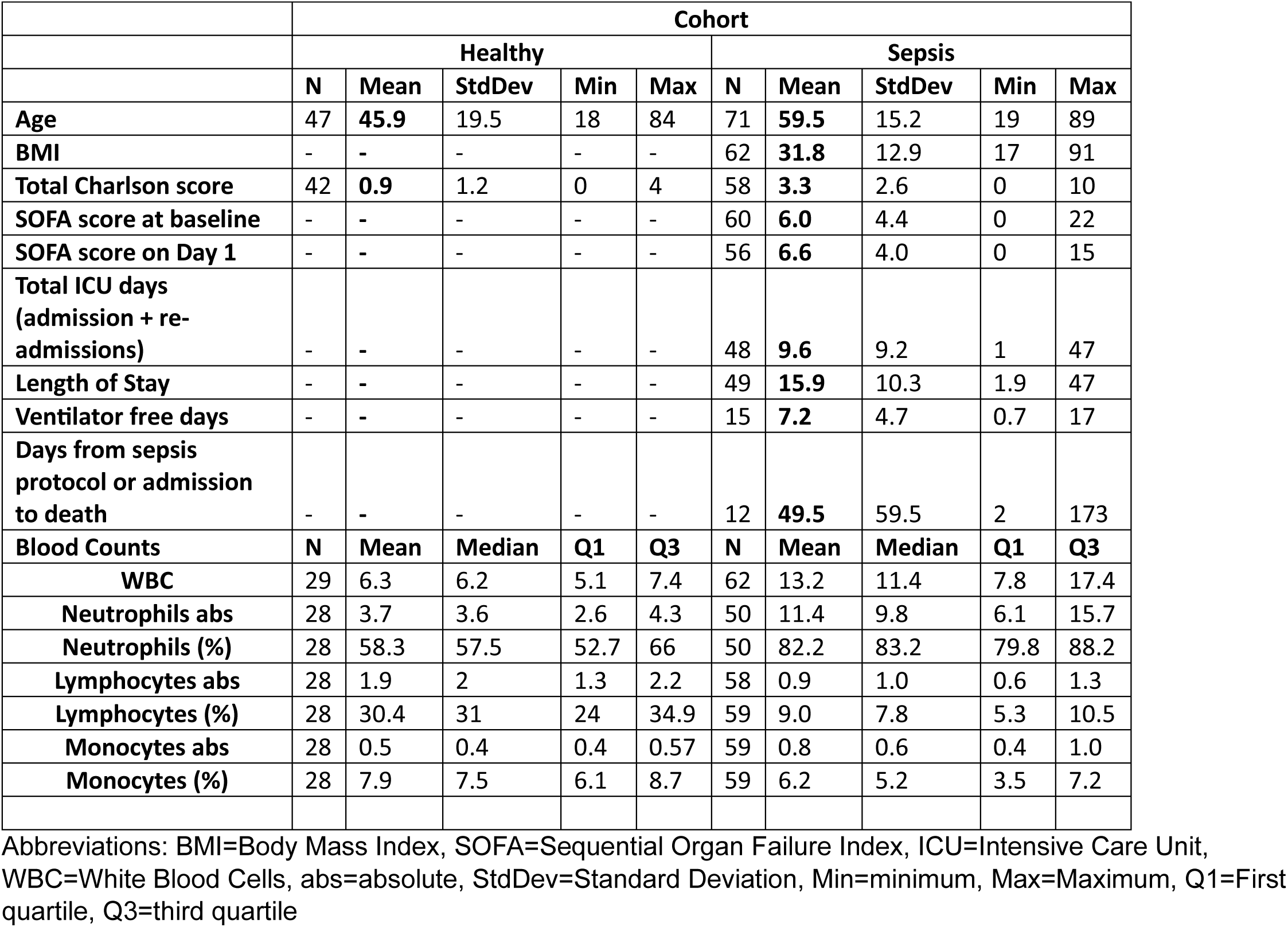
Patient Clinical Characteristics.

**Table 2:** Patient Demographics and Disposition.

|  | Cohort |  |  |  |
| --- | --- | --- | --- | --- |
|  | Healthy |  | Sepsis |  |
|  | N= |  |  |  |
|  | 49 | % | 78 | % |
| **Sex (n=149)** |  |  |  |  |
| Male | 21 | (50) | 38 | (64.5) |
| Female | 26 | (44.3) | 35 | (29) |
| **Age Group (n=147)** |  |  |  |  |
| Under 50 | 26 | (53.1) | 21 | (26.9) |
| 50 - 59 | 8 | (16.3) | 14 | (17.9) |
| 60 - 69 | 6 | (12.2) | 15 | (19.2) |
| 70 - 80 | 6 | (12.2) | 15 | (19.2) |
| Over 80 | 1 | (2) | 6 | (7.7) |
| **Secondary infections (n=40)** |  |  |  |  |
| **Disposition at discharge (n=78)** |  |  |  |  |
| Home |  |  | 9 | (11.5) |
| Home with service |  |  | 14 | (17.9) |
| Inpatient rehabilitation facility |  |  | 10 | (12.8) |
| Skilled nursing facility |  |  | 5 | (6.4) |
| Long term acute care |  |  | 2 | (2.6) |
| Hospital |  |  | 1 | (1.3) |
| Hospice |  |  | 2 | (2.6) |
| Death |  |  | 9 | (11.5) |
| Against medical advice |  |  | 1 | (1.3) |
| **Disposition at follow-up (n=17)** |  |  |  |  |
| Home |  |  | 7 | (9) |
| Skilled nursing facility |  |  |  |  |
| Hospital |  |  | 2 | (2.6) |
| Death |  |  | 4 | (5.1) |

*Exclusion criteria*: Individuals with autoimmune diseases being treated with biologic immune modulators were excluded, as were individuals who had received antineoplastic therapies or those diagnosed with cancer within the previous six months. Vulnerable populations were also excluded.

Samples for ELISpot were collected from sepsis patients within the first 72 hours of sepsis diagnosis. Additional blood was collected concurrently for complete blood count analysis by the clinical laboratories at each institution.

*Study approval*. Centralized ethics approval was obtained from the University of Florida Institutional Review Board (IRB 202000924), which served as the sponsoring institution for this multicenter clinical study. Written informed consent was obtained from each patient or their proxy decision maker at individual clinical sites.

### ELISpot

ELISpot was performed as previously described (7, 11). Briefly, ELISpot specific for IFN-γ and TNF-α was carried out using diluted whole blood in CTL-Test™ media (Cellular Technologies Ltd, CTL, Cleveland, OH, USA) supplemented with 2mM L-Glutamine (ThermoFisher Scientific, Waltham, MA, USA). Immunomodulatory agents included LPS (ENZO Life Sciences, Farmingdale, NY, USA), anti-CD3 and anti-CD28 mAb (BioLegend, S Diego, CA, USA), recombinant human IL-7 (R&D Systems, Minneapolis, MN, USA), anti-PD1 mAb (Nivolumab, Bristol Myers Squibb, New York, NY, USA), Dexamethasone (Viatris, formerly Mylan, Canonsburg, PA, USA), and Resiquimod (Invivogen, San Diego, CA, USA), using IFN-γ and TNF-α ELISpot kits (CTL). IFN-γ and TNF-α ELISpots were incubated for 22 hours before washing with PBS+0.05% Tween (MilliporeSigma) and developing per manufacturer instructions (CTL). Concentrations of stimulants and adjuvants used are listed in **Supplemental Table 1**.

ELISpots were analyzed using CTL S6 Entry ELISpot readers at each site with ImmunoSpot 7.0 professional software (CTL). Instruments across all sites were calibrated by CTL to minimize instrument-to-instrument variability. Results are presented as Spot Forming Units (SFU), which represents the number of cells in the assay which produce cytokine, Spot Size (SS, expressed in µm^2^), a measure of the mean diameter of spots in the assay, and Total Well Intensity (TWI), a measure which uses the proportion of the well which is covered by spots, and intensity of coverage, to provide a measure comparable to the results obtained by ELISA (7).

### Statistical Analysis

ELISpot were performed in duplicate and the results per sample averaged. Data were analyzed using Graphpad Prism 10.1.2 (Graphpad, San Diego, CA, USA). Analysis of differences between groups were performed using a paired, non-parametric Friedman Test with multiple comparisons, or Wilcoxon Test as appropriate. Significant comparisons are indicated on figures, non-significant comparisons are not displayed.

## RESULTS

### IL-7 but not anti-PD-1 mAb increases IFN-γ production in whole blood ELISpot

Recombinant human Interleukin-7 (IL7) and anti-Programmed cell Death protein-1 (anti-PD1) mAb (Nivolumab) were tested in a diluted whole blood ELISpot for their ability to enhance lymphocyte IFN-γ production above the baseline production with anti-CD3/CD28 mAb stimulation. IL-7 induced an ∼ 85% (p<0.0001) increase in the number of IFN-γ spot forming units (SFU) in healthy control subjects, whereas there was a more modest increase of ∼37% (p<0.0001) in the number of IFN-γ SFUs, in blood from septic patients (**Fig. 1**, **Table 3).** The relative amount of IFN-γ produced per lymphocyte is indicated by the spot size (SS). IL-7 caused an increase of 17% (p<0.0001) and 11% (p<0.0001) in spot size in healthy subjects and septic patients, respectively. IL-7 also increased the total well intensity (TWI), a measurement that reflects both the area of the well positive for cytokine production and the colorimetric intensity of the coverage, in healthy subjects and septic patients by 127% (p<0.0001) and 79% (p<0.0001), respectively. Importantly, the effect of IL-7 to increase IFN-γ production occurred in >90% of both healthy control subjects and patients with sepsis (**Fig. 2**).

**Figure 1:**
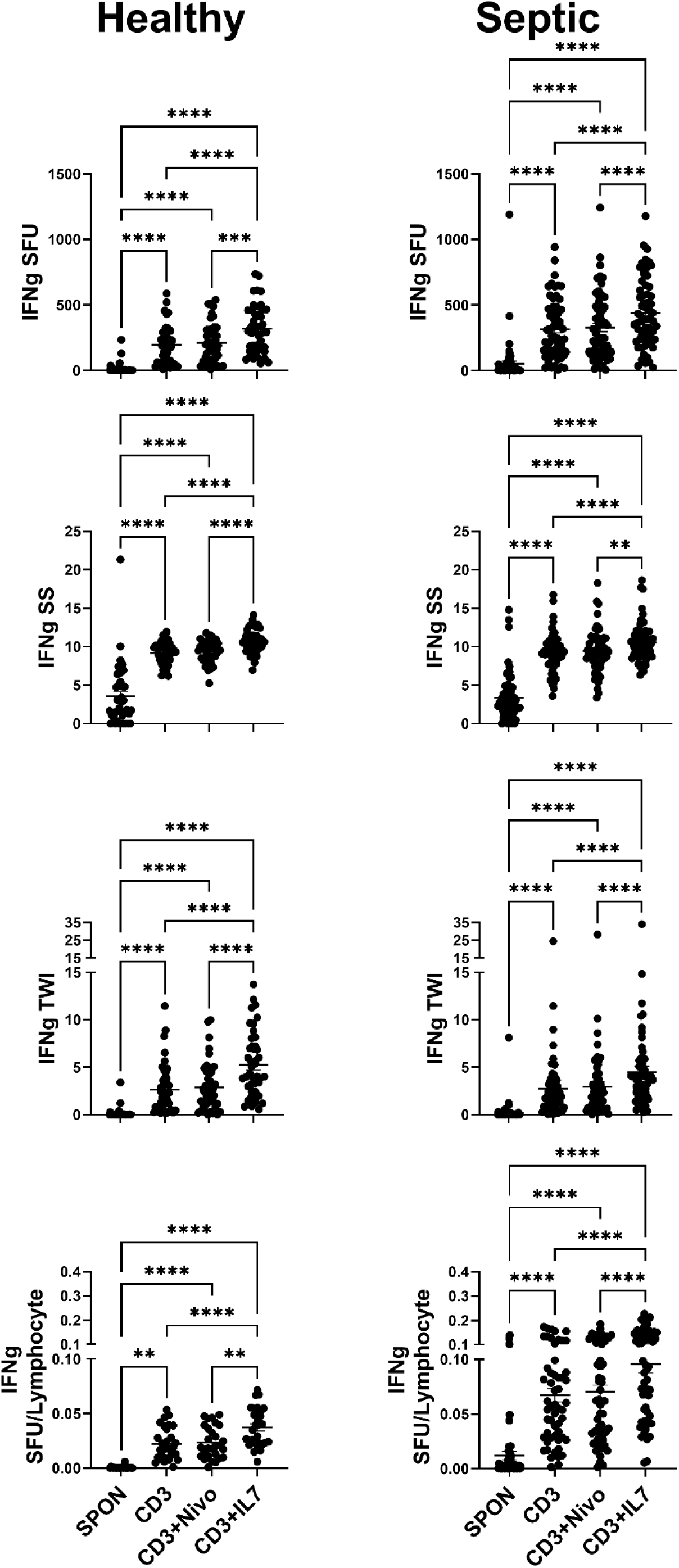
Adjuvant effect on IFN-γ production in conjunction with anti-CD3/CD28 mAb stimulation. Samples were stimulated with anti-CD3/28mAb with and without either anti-PD-1 mAb (Nivolumab; NIVO) or IL-7. Anti-CD/CD28 mAb (CD3) increased IFN-γ SFU (upper panels), SS (upper-middle panels) and TWI (lower-middle panels) in samples from both Healthy donors (left, n=48) and Septic patients (right, n=61) as compared to unstimulated spontaneous (SPON) production. NIVO did not increase CD3 induced IFN-γ SFU, SS, or TWI, while IL-7 did increase CD3 induced IFN-γ SFU, SS, and TWI. SFU (Top panels) were normalized to the number of lymphocytes plated in the ELISpot well (Bottom panels). Lymphocyte number per well was determined by multiplying the draw specific ALC (K/cumm) by 1000 (converting K/cumm to cells/cumm) then by the volume of blood added to the well (5µL). Pairwise statistical relationships of SFU/Lymphocyte remain largely the same as SFU. *p<0.05, **p<0.01, ***p<0.001, ****p<0.0001

**Figure 2:**
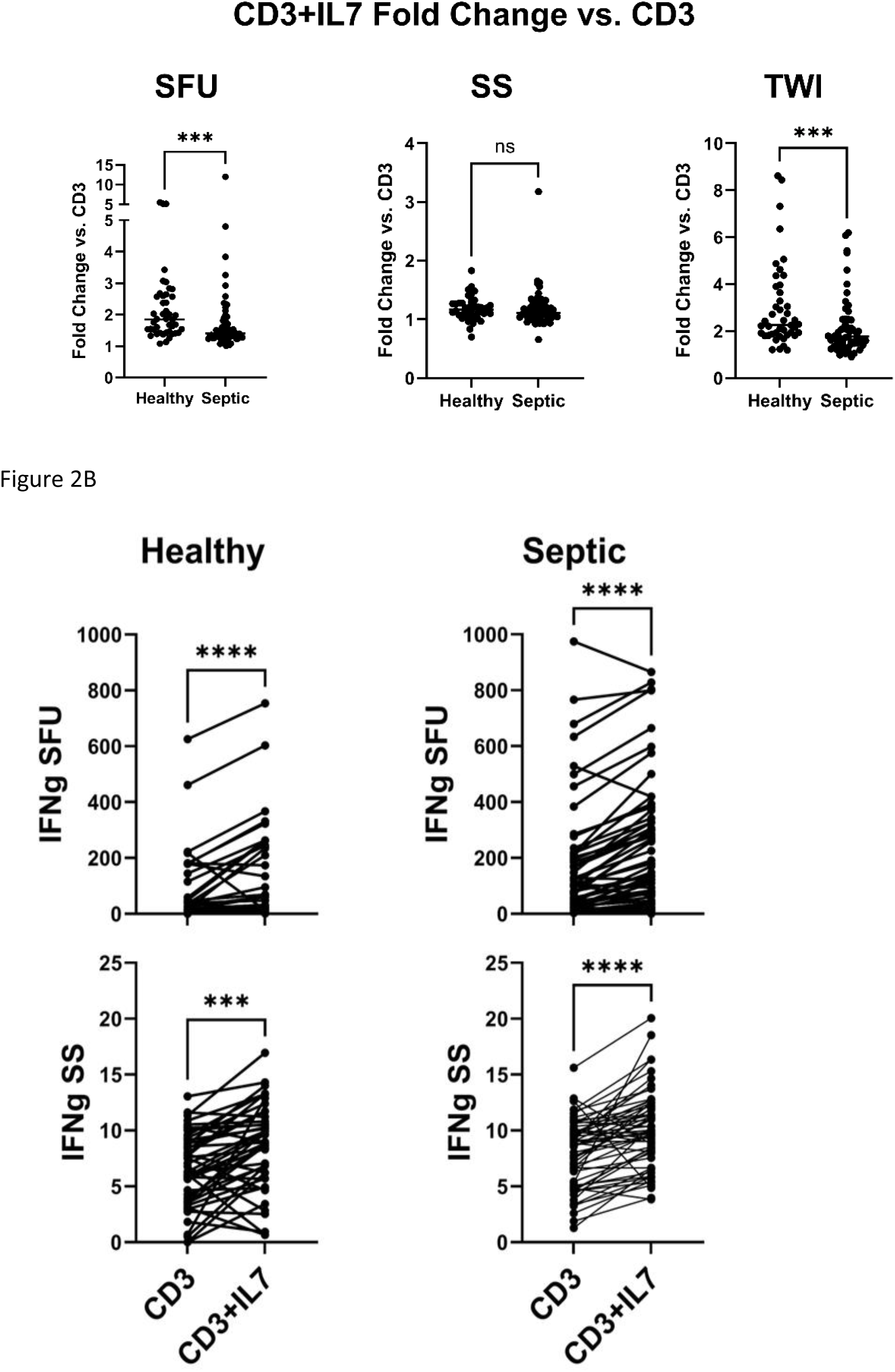
Difference in relative IFN-γ response to IL-7 between healthy donors and septic patients. **A**. IL-7 fold-changes were calculated for SFU, SS, and TWI for all samples; anti-CD3/28 mAb (CD3)+IL-7 values were divided by CD3 alone to yield fold-change. Samples from healthy donors displayed a higher responsiveness to IL-7 in the SFU (Left) and TWI (Right) measurements, based on fold-change, than did those from septic patients. **B.** Visualization of paired data points for CD3 alone or with IL-7 for samples from healthy donors (Left, n=48) and septic patients (Right, n=61). The bulk of samples show an increase of SFU and SS with IL-7 treatment *p<0.05, **p<0.01, ***p<0.001, ****p<0.0001.

**Table 3:** IFN-γ ELISpot Data.

|  | SPON |  | CD3 |  | CD3+IL-7 |  | CD3+NIVO |  |
| --- | --- | --- | --- | --- | --- | --- | --- | --- |
|  | Healthy | Septic | Healthy | Septic | Healthy | Septic | Healthy | Septic |
| SFU | 2<br>(0.88,4) | 7<br>(1,30.5) | 182.5<br>(57.5,282.88) | 255<br>(142,470) | 296.5<br>(161.75,458.25) | 389<br>(239.5,612) | 193.5<br>(80.38,309.88) | 253<br>(143,485) |
| SS | 2.43<br>(1.08,5.59) | 2.46<br>(1.63,5.229) | 9.54<br>(8.28,10.23) | 9.25<br>(7.88,10.42) | 10.64<br>(9.88,11.46) | 10.36<br>(9.05,11.53) | 9.77<br>(8.47,10.49) | 9.44<br>(8.20,10.93) |
| TWI | 0.01<br>(0.00,0.02) | 0.01<br>(0.001,0.1) | 2.07<br>(0.64,3.68) | 1.92<br>(0.86,3.35) | 4.19<br>(2.45,7.17) | 3.69<br>(1.89,5.08) | 2.31<br>(0.95,4.5) | 2.03<br>(0.67,3.75) |
| SFU<br>FC |  |  |  |  | 1.85<br>(1.45,2.58) | 1.37<br>(1.26,1.80) | 1.10<br>(1.01,1.20) | 1.06<br>(0.96,1.15) |
| SS<br>FC |  |  |  |  | 1.17<br>(1.07,1.27) | 1.11<br>(1.04,1.23) | 1.03<br>(0.96,1.07) | 1.02<br>(0.94,1.10) |
| TWI<br>FC |  |  |  |  | 2.27<br>(1.87,3.71) | 1.79<br>(1.47,2.30) | 1.11<br>(0.99,1.33) | 1.09<br>(0.94,1.24) |

All values are "Median (Quartile 1, Quartile 3)". Abbreviations: SFU=Spot Forming Units, SS=Spot Size, TWI=Total Well Intensity, FC=Fold change (relative to CD3)

Lymphopenia is a hallmark of patients with sepsis due to the profound sepsis-induced apoptosis, and this decrease in lymphocytes in septic patients compared to healthy subjects was present in the current study (**Table 1**). To examine the impact of this difference in the number of lymphocytes in septic vs healthy volunteers, the data were normalized by dividing the spot number by the total number of lymphocytes plated in each well. Lymphocyte number per well was determined by multiplying the absolute lymphocyte count (ALC) specific to each blood sample (K/mm^3^) by 1,000 (converting K/mm^3^ to cells/mm^3^) then by the volume of blood added to the well (5 µL) [=ALC*1000*5]. This normalization protocol allowed for the ELISpot results to be presented as SFU per number of lymphocytes present in the ELISpot well, (SFU/Lymphocyte, **Fig. 1**). Normalization of the SFU, to adjust for the actual number of lymphocytes plated in each well, showed the percentage of lymphocytes producing IFN-γ was higher in septic patients compared to healthy subjects (p<0.0001, **Fig. 3**), similar to previous findings (7).

**Figure 3:**
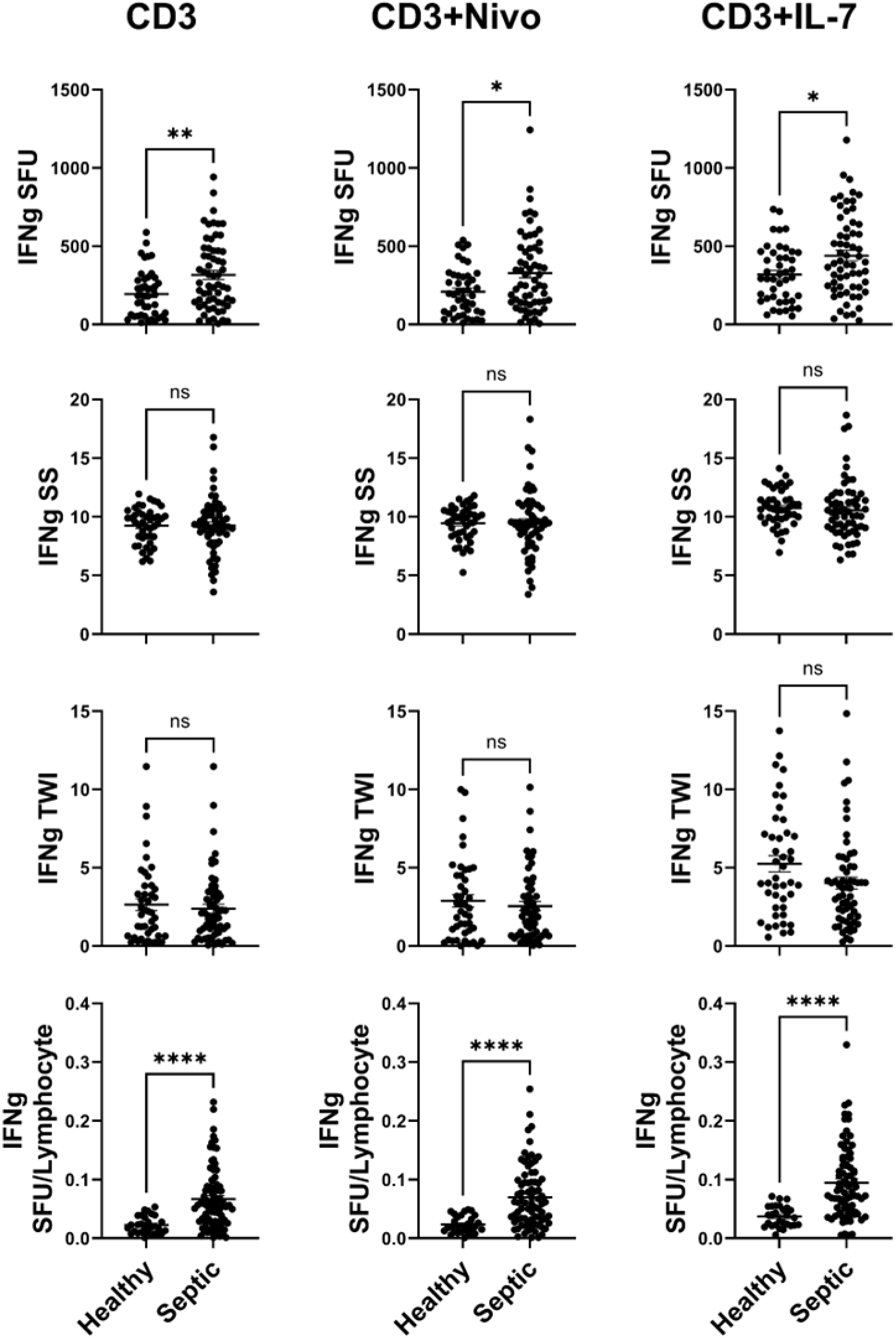
Septic patient samples produce greater IFN-γ SFU and SFU/Lymphocyte than Healthy donor samples. Comparison between Healthy donor (n=48) and Septic patient (n=61) whole blood samples shows septic blood samples have higher IFN-γ SFU (top) than healthy donor samples in response to anti-CD3/28 mAb (CD3) alone (left), CD3+anti-PD-1 mAb (Nivolumab; NIVO) (middle), and CD3+IL-7 (right). To account for high variability in cell counts among septic patients and between septic and healthy samples, SFU were normalized to number of lymphocytes plated in ELISpot wells (bottom). When SFU were normalized to lymphocyte numbers, the differences between septic and healthy IFN-γ production became more apparent. As no significant differences were discerned between CD3 treated samples and CD3+NIVO treated samples, the differences displayed in the “CD3+NIVO” column between healthy and septic are a consequence of the greater responsiveness of septic samples to CD3 treatment. *p<0.05, **p<0.01, ***p<0.001, ****p<0.0001.

In contrast to IL-7, anti-PD-1 mAb failed to increase IFN-γ production in either healthy control or septic patients by any parameter, i.e., SFU, SS, or TWI (**Fig. 1**). Because of concern that baseline anti-CD3/ CD28 mAb stimulation may have masked the effect of anti-PD-1 mAb to increase IFN-γ, additional studies using a lower concentration of anti-CD3/ CD28 were performed. Despite using the lower doses of anti-CD3/ CD28, anti-PD-1 mAb failed to increase either the IFN-γ spot number or size (**Supplemental Figs. 1-3**).

### Dexamethasone reduces LPS and resiquimod-induced TNF-α production

Corticosteroids have been widely used therapeutically in patients with sepsis to dampen the hyper-inflammatory response seeking to minimize pro-inflammatory cytokine mediated tissue damage. Dexamethasone (DEX) is a potent corticosteroid that demonstrated survival benefits in patients with COVID-19 (12). We tested the effect of DEX on TNF-α production at a concentration of DEX that reflected the circulating blood level in patients being treated with DEX at a total daily dose of 6 mg, a commonly used therapeutic dose for COVID-19 patients (12). Diluted whole blood samples were plated either with LPS or resiquimod (R848), as primary stimulants. Both R848 and LPS potently stimulated TNF-α production and increased SFU, SS, and TWI (**Figs. 4-5**). DEX significantly decreased both spontaneous and TLR agonist-stimulated TNF-α production. DEX reduced R848 induced SFU, SS, and TWI by approximately 35% (p<0.01), 18% (p<0.0001), and 54% (p<0.0001), respectively in healthy donor samples and 36% (p<0.0001), 20% (p<0.05), and 61% (p<0.0001) in samples from septic patients. Meanwhile, DEX reduced LPS induced SFU, SS, and TWI by approximately ∼37% (p<0.0001), 31% (p<0.0001), and ∼61% (p<0.0001) respectively in healthy donor samples and ∼42% (p<0.0001), 25% (p<0.0001), and 64% (p<0.0001) respectively in sepsis samples. Even in the presence of DEX, the TLR-agonist-stimulated TNF-α SFU and TWI was still significantly greater than spontaneous production (**Figs 4-5**, **Table 4**).

**Figure 4:**
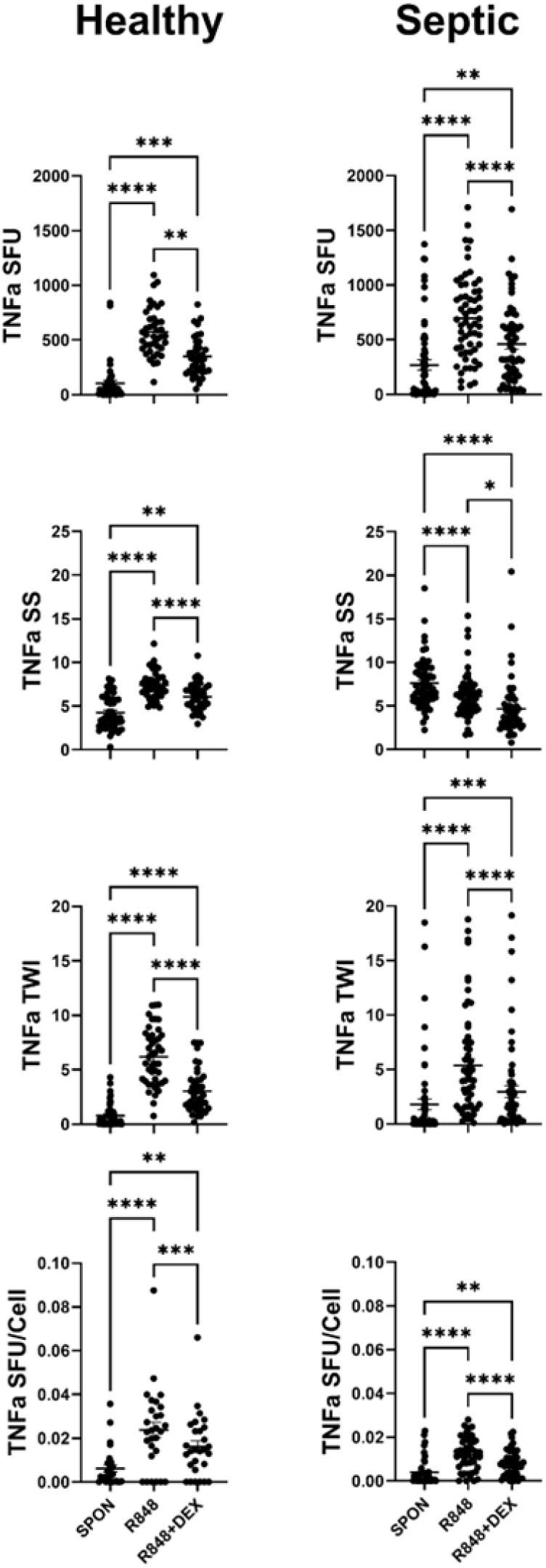
Adjuvant effect on TNF-α production in conjunction with resiquimod (R848) stimulation. Samples were stimulated with R848 with and without desamethasone (DEX). R848 increased TNF-α SFU (upper panels), SS (upper-middle panels) and TWI (lower-middle panels) in samples from both Healthy donors (left, n=48) and Septic patients (right, n=61) as compared to unstimulated (spontaneous, SPON) production. DEX markedly abrogated R848 induced TNF-α production in SFU, SS, and TWI. SFU (Top panels) were normalized to the number of TNF-α producing cells (Cells) plated in the ELISpot well (Bottom panels). The number of TNF-α producing cells per well was determined by multiplying the draw specific Absolute Neutrophil and Absolute Monocyte counts (ANC+AMC) in K/cumm by 1000 (converting K/cumm to cells/cumm) then by the volume of blood added to the well (5µL). Pairwise statistical relationships of SFU/Cell remain largely the same as SFU. *p<0.05, **p<0.01, ***p<0.001, ****p<0.0001.

**Figure 5:**
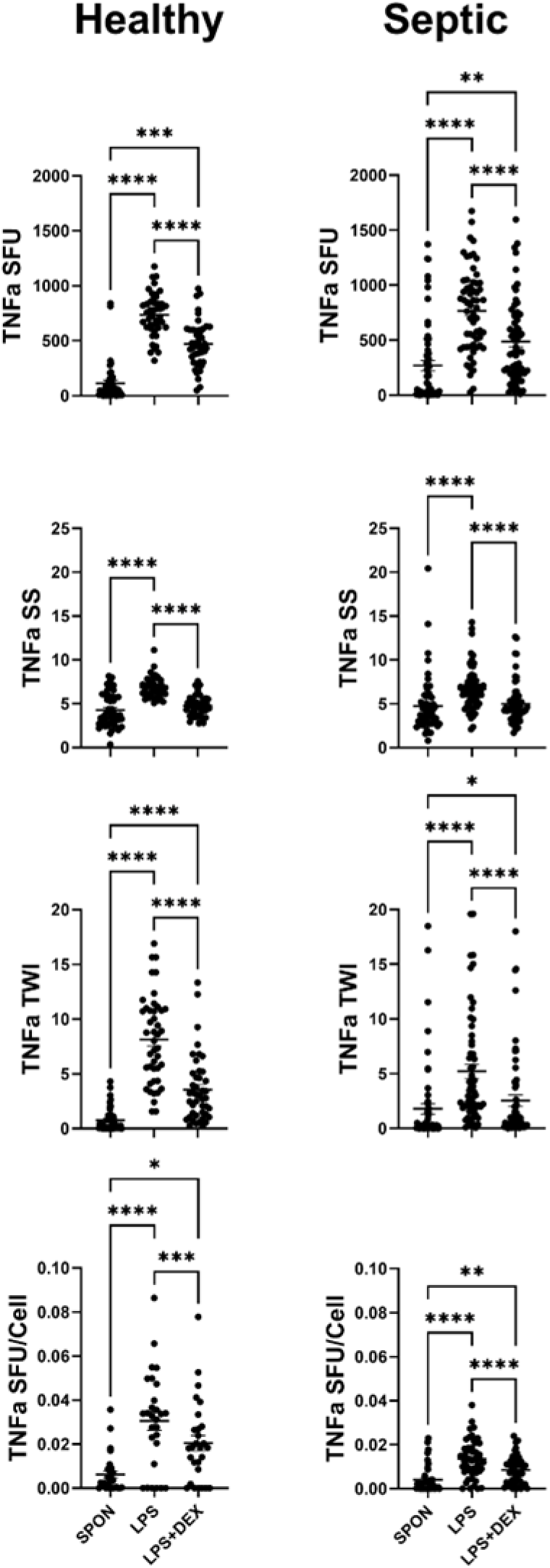
Adjuvant effect on TNF-α production in conjunction with Lipo-polysaccharide (LPS) stimulation. Samples were stimulated with LPS with and without dexamethasone (DEX). LPS increased TNF-α SFU (upper panels), SS (upper-middle panels) and TWI (lower-middle panels) in samples from both Healthy donors (left, n=48) and Septic patients (right, n=61) as compared to unstimulated (spontaneous, SPON) production. DEX markedly abrogated LPS induced TNF-α production in SFU, SS, and TWI. SFU (Top panels) were normalized to the number of TNF-α producing cells (Cells) plated in the ELISpot well (Bottom panels). The number of TNF-α producing cells per well was determined by multiplying the draw specific Absolute Neutrophil and Absolute Monocyte counts (ANC+AMC) in K/cumm by 1000 (converting K/cumm to cells/cumm) then by the volume of blood added to the well (5µL). Pairwise statistical relationships of SFU/Cell remain largely the same as SFU *p<0.05, **p<0.01, ***p<0.001, ****p<0.0001.

**Table 4:** TNF-α ELISpot Data.

|  | Spon |  | LPS |  | LPS+Dex |  | R848 |  | R848+Dex |  |
| --- | --- | --- | --- | --- | --- | --- | --- | --- | --- | --- |
|  | Healthy | Septic | Healthy | Septic | Healthy | Septic | Healthy | Septic | Healthy | Septic |
| SFU | 56.5<br>(12.75,<br>129.25) | 100.5<br>(15.0,<br>363.5) | 741.5<br>(621.38,<br>846.63) | 790.5<br>(450.5,<br>1017) | 448.5<br>(306.25,<br>599.5) | 397.5<br>(199,<br>721.5) | 532.5<br>(425.13,<br>671.38) | 661<br>(410.5,<br>939) | 330.5<br>(219.25,<br>486.13) | 334.5<br>(176.5,<br>623.5) |
| SS | 3.64<br>(2.72, 5.84) | 4.00<br>(2.83, 5.75) | 6.66<br>(6.08, 7.45) | 6.53<br>(5.02, 7.63) | 4.75<br>(3.91, 5.33) | 4.3<br>(3.53, 5.48) | 7.24<br>(6.40, 8.08) | 7.18<br>(5.81, 9.34) | 6<br>(5.02, 6.95) | 5.99<br>(4.47, 7.02) |
| TWI | 0.31<br>(2.72, 5.84) | 0.21<br>(0.02, 1.66) | 8.61<br>(5.2, 10.82) | 3.46<br>(1.88, 7.62) | 2.92<br>(1.285, 4.77) | 0.83<br>(0.25, 3.74) | 5.59<br>(3.97, 8.18) | 4.05<br>(1.58, 7.47) | 2.61<br>(1.75, 4.07) | 1.60<br>(0.3, 3.65) |
| SFU<br>FC |  |  |  |  | 0.64<br>(0.53, 0.76) | 0.59<br>(0.41, 0.76) |  |  | 0.65<br>(0.51, 0.75) | 0.64<br>(0.46, 0.72) |
| SS<br>FC |  |  |  |  | 0.68<br>(0.63, 0.77) | 0.71<br>(0.60, 0.82) |  |  | 0.82<br>(0.77, 0.87) | 0.80<br>(0.74, 0.87) |
| TWI<br>FC |  |  |  |  | 0.37<br>(0.27, 0.47) | 0.29<br>(0.17, 0.47) |  |  | 0.46<br>(0.37, 0.53) | 0.39<br>(0.24, 0.52) |

All values are "Median (Quartile 1, Quartile 3). Abbreviations: SFU=Spot Forming Units, SS=Spot Size, TWI=Total Well Intensity, FC=Fold change (relative to either LPS or R848 as appropriate)

As previously reported by our group, whole blood stimulated TNF-α production is due not only to monocytes but also neutrophils which can make surprisingly large amounts of the cytokine (7). The absolute number and proportion of monocytes and neutrophils in septic patients is frequently different than in healthy subjects because of the active host inflammatory response occurring during an acute infection. Consequently, normalization of the ELISpot results was performed by dividing number of TNF-α producing cells (SFUs) by the total number of TNF-a producing cells plated in the well to account for the varying numbers of TNF-α producing cells in the individual sample. The number of TNF-α producing cells per well was determined by multiplying the Absolute Neutrophil (ANC) and Absolute Monocyte (AMC) counts in each patient blood sample in K/cumm by 1000 (converting K/cumm to cells/cumm) then by the volume of blood added to the well (5µL) [=(ANC+AMC)*1000*5]. After normalization, the relationship and overall impact of DEX to decrease TNF-α production remained consistent with non-normalized spot forming units for both LPS and resiquimod stimulation (**Fig. 6**).

**Figure 6:**
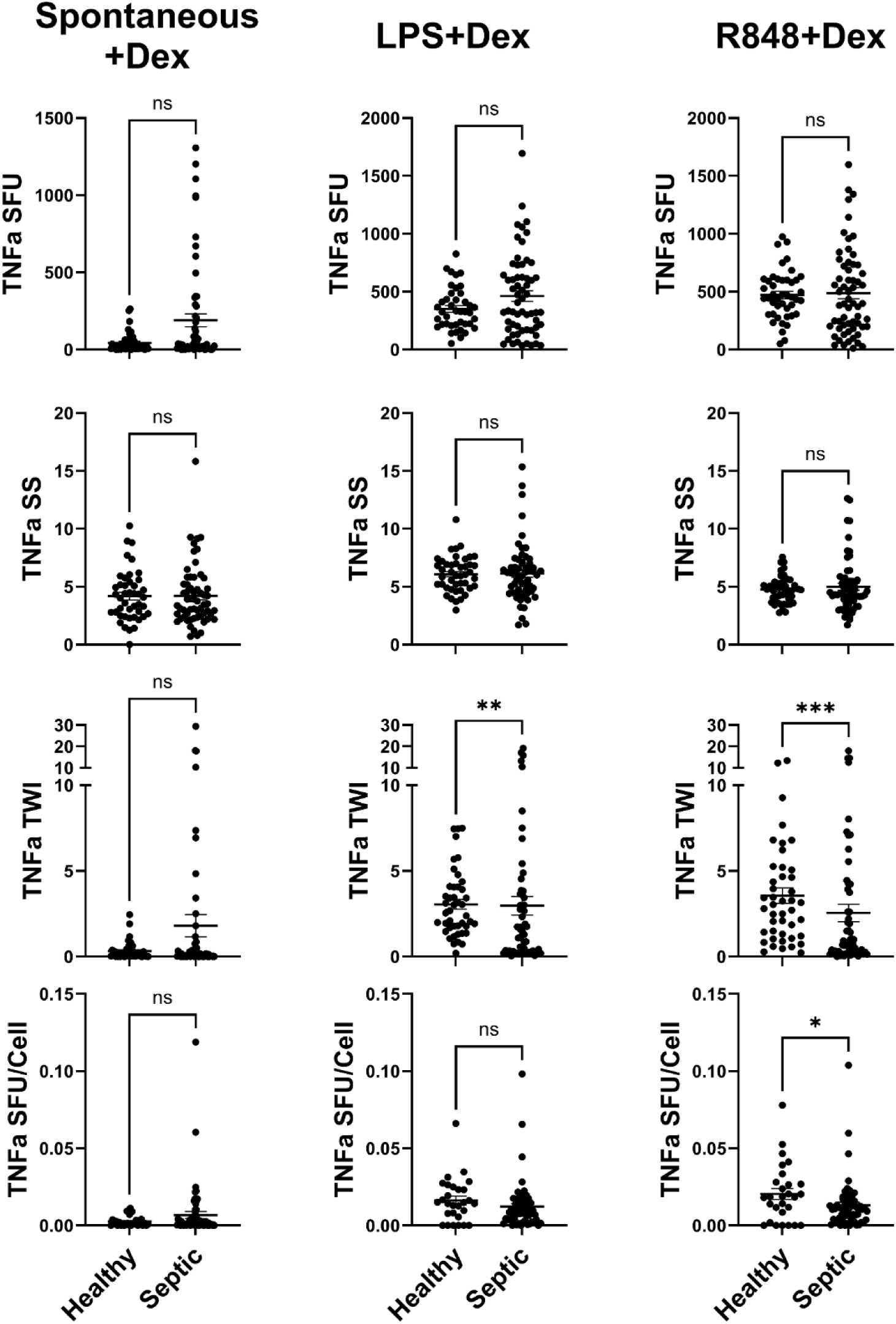
Comparison of responsiveness to dexamethasone between healthy donor and septic samples. Dexamethasone (DEX) was able to abrogate TNF-α production equally in healthy donor (n=48) and septic patient (n=61) samples for most parameters when looking at spontaneous TNF-α production (Left), LPS-induced TNF-α production (middle), or R848-induced TNF-α production (Right). DEX-based inhibition of LPS-induced TNF-α production was more pronounced in septic samples in the TWI measurement and R848-induced TNF-α production on the TWI measurement and on an SFU per cell basis in septic patient than in healthy donor samples. *p<0.05, **p<0.01, ***p<0.001, ****p<0.0001.

### Dexamethasone reduces TNF-α comparably in healthy subjects and septic patients

Comparison of the effect of DEX on stimulated TNF-α production in healthy subjects versus septic patients showed that the magnitude of TNF-α suppression was comparable (**Fig. 6**). Spontaneous and stimulated TNF-α production were largely similar in healthy subjects and septic patients (**Supplemental Fig. 4**), and DEX reduced TNF-α SFU and SS to nearly equal extents. However, DEX reduced TWI of LPS and R848 to a greater extent in septic patients versus healthy subjects (p<0.01 and p<0.001, respectively). Septic SFU/cell was also more greatly reduced than healthy in the latter case (p<0.05).

### IL-7 reduces the suppressive effect of dexamethasone on stimulated IFN-γ production

In addition to examining the effect of DEX to reduce TNF-α production, we also tested the impact of DEX on lymphocyte IFN-γ production. DEX had a potent suppressive effect on anti-CD3/ CD28 mAb stimulated IFN-γ production in both healthy subjects and in septic patients (**Fig. 7**). DEX decreased IFN-γ SFU in healthy subjects and septic patients by ∼51% (p<0.01) and ∼37% (p<0.0001) respectively. Similarly, DEX decreased IFN-γ SS by ∼12% (p<0.001) in healthy subjects but not in septic patients (**Fig. 7**). Importantly, DEX also diminished but did not eliminate the positive effect of IL-7 to increase lymphocyte IFN-γ production. Although DEX reduced SFU and SS, the DEX-induced reduction in IFN-γ in the IL-7 treated healthy control and septic patient samples remained significantly greater than in the non-IL-7 treated samples (p < 0.001 for SFU, SS, and TWI in healthy, p<0.0001 for SFU and TWI in septic). Thus, IL-7 was able to partially overcome the suppressive effect of DEX on IFN-γ production.

**Figure 7:**
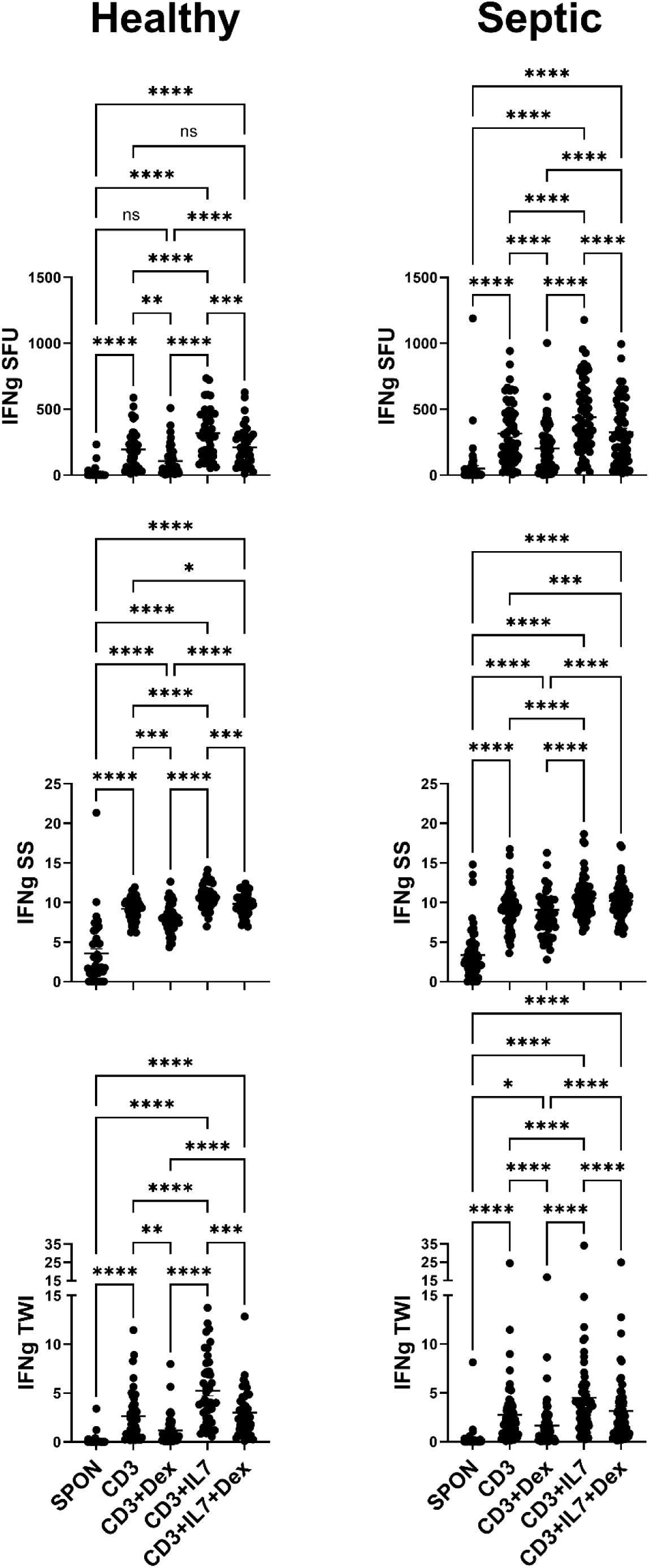
Effect of dexamethasone on IFN-γ production. IFN-γ SFU (Top), SS (Middle), and TWI (Bottom), are reduced when desamethasone (DEX) is added to ELISpot. DEX abrogates anti-CD3/28 mAb (CD3) induced IFN-γ production. DEX-induced reduction of IFN-γ production is still present when co-cultured with CD3 and IL-7, though there is still an IL-7 effect present even with DEX co-treatment. Results were similar in healthy donor (n=48) and septic patient (n=61) samples. *p<0.05, **p<0.01, ***p<0.001, ****p<0.0001.

### IFN-γ production is increased in septic patients vs healthy subjects

Comparison of IFN-γ production in healthy subjects versus septic patients showed septic patients had an increase in IFN-γ SFU and SFU/Lymphocyte compared with healthy subjects, and this difference was significant for stimulated whole blood with and without anti-PD-1 mAb, or anti-CD3/28 mAb + IL-7 (**Fig. 3**). However, there was no difference in either SS nor TWI between septic patients and healthy subjects for any condition. The increase in the number of IFN-γ producing lymphocytes that occurred in septic patients versus healthy donors in the anti-CD3/ CD28 mAb stimulated samples also occurred in the IL-7 and anti-PD-1 mAb treated samples (**Fig. 3**). Since there is no difference between anti-CD3/ CD28 mAb alone and with anti-PD-1 mAb, the difference is likely to be a consequence of the anti-CD3/CD28 mAb effect alone rather than an effect of anti-PD-1 mAb.

The magnitude of change between healthy subjects and septic patient samples varied significantly; Despite higher SFU and SFU/lymphocyte, the fold change of SS and TWI induced by IL-7 co-stimulation trended lower for the septic patients than healthy subjects (**Fig. 3**), though this difference did not reach the level of significance.

When the dose of anti-CD3/CD28 mAb stimulation was reduced, the relationship remained consistent. SFU, TWI, and SFU/Lymphocyte were consistently higher in the septic group with IL-7 and anti-PD-1 mAb co-stimulation (**Supplemental Fig. 2**). However, there was no difference in the degree of change between the healthy subject and septic patient groups for low dose anti-CD3/28 mAb stimulation (**Supplemental Figure 3**).

## DISCUSSION

Sepsis initiates a diverse immune response that varies significantly and is dependent upon both the stimuli and any pre-existing comorbidities (13–15). Furthermore, there is often a dissociation of the innate immune response which consists of early activation and the adaptive immune response which is frequently impaired. Methods to endotype the immune response have also been complicated by the fact that the patients’ immune responses change over time depending on clinical trajectory and outcome. Previous work by our group has shown that ELISpot can distinguish between the innate and adaptive immune responses to sepsis by using LPS-stimulated TNF-α to reflect innate immunity and anti-CD3/CD28 mAb stimulation to induce IFN-γ to reflect adaptive immunity (7, 16, 17). Septic patients who had poor outcomes had an association with impaired IFN-γ response as indicated by a decrease in both the number of IFN-γ producing lymphocytes and amount of IFN-γ produced per lymphocyte (16,17). Furthermore, recent studies from our team have shown that septic patients who have an increased *ex vivo* whole blood TNF-α response have an increased survival probabilities compared with septic patients who made suppressed TNF-α production (*manuscript submitted*). Thus, ELISpot joins the growing list of methods to identify patients who are at high risk of dying of sepsis.

While information of likely patient outcomes is useful, knowledge of the *functional state* of the patients’ immune system (i.e. whether the patient is in a hyper-inflammatory or immunosuppressed state) would enable effective application of drug therapies to restore immune homeostasis and improve morbidity and mortality in sepsis. Results of the present study indicate that ELISpot can not only provide an assessment of the functional state of the patient’s host immunity, but it can also help identify specific immune modulatory drugs that are most likely to be effective in restoring immune homeostasis in specific patients. The potential importance of developing a test to identify the optimal drug therapy to restore patient immune homeostasis could have an enormous impact on sepsis outcomes. Many investigators have argued that the failure of the various immune drug therapy trials in sepsis stems from the drugs being given to patients indiscriminately without knowledge of the underlying immune-pathobiology and endotypes. (5,7). For example, studies indicate that administration of corticosteroids to patients in refractory septic shock is likely to be beneficial in the subset of patients who are in the more hyper-inflammatory phase but may be detrimental to those patients who are in the immunosuppressive phase of the disorder (5,6). Recently, investigators have argued that an evaluation of the septic patients’ immune status is a “*necessary precondition*” for conducting trials of immune modulatory drug therapies (5).

An ideal test to help guide immune modulatory drug therapies in sepsis would not only provide information about the overall status of the patient’s immune function but also concurrently identify which of the various immune modulatory drugs would be most effective in restoring the patients’ immune homeostasis (7). Because of the remarkable success of checkpoint inhibitors (e.g. nivolumab or ipilimumab) in oncology, there is a burgeoning list of drugs currently available and in development that effectively modulate the cellular elements of the innate and adaptive immune system. In this regard, recent work by our group using a clinically relevant murine model of sepsis showed ELISpot tracked the dynamic changes in the innate and adaptive immune system progressing from the early hyper-inflammatory phase to the later immune recovery state (18). Importantly, ELISpot also detected the expected changes in adaptive and innate immunity occurring in mouse circulating white blood cells and in splenic immune effector cells *in response to treatment* with corticosteroids and other immune modulatory drug therapies (18). The results of the present study indicate ELISpot is also capable of: 1) evaluating the effectiveness of immune modulatory drug therapies *ex vivo* in fresh whole blood samples from patients with sepsis, 2) displaying the ability of drugs to independently regulate the innate and adaptive host response to sepsis, and 3) elucidating the net effect of drugs with competing actions on host immunity. For example, IL-7 partially reversed the immunosuppressive effect of corticosteroids on lymphocytes (**Fig 7**).

There are several interesting findings from the present study that have potential clinical implications. The effect of IL-7 to improve lymphocyte IFN-γ production in >90% of samples from patients with sepsis suggests IL-7 could be effective in reversing immune suppression in septic patients who are in the immunosuppressive phase. Although the current study is an *ex vivo* study of patient blood, septic mice treated *in vivo* with IL-7 show comparable increases in lymphocyte IFN-γ production (18). Additionally, two case reports of patients with intractable life-threatening infections which were not responding to standard antimicrobial therapies showed a marked improvement in ELISpot IFN-γ production after initiation of IL-7 therapy (19, 20). These changes were associated with improved survival as well. Although the present studies only examined the effect of IL-7 on lymphocyte IFN-γ production, additional studies from our group showed IL-7 improved lymphocyte polyfunctionality by acting to increase lymphocyte production of numerous other cytokines including TNFα, IL-1β, IL-6, IL-17A, and IL-10 (T.S. Griffith, unpublished data). A recent report by Tilsed and colleagues provides a potential explanation for the impressive ability of IL-7 to increase IFN-γ in almost all healthy control and septic patient blood samples (21). These investigators performed transcriptomic analysis of splenic murine CD8 T cells treated with IL-7 and showed that IL-7 but not IL-2 or IL-15 (other members of the common γ-chain cytokine family) markedly enhanced genomic pathways associated with protein expression. This finding likely also explains why IL-7 increase lymphocyte polyfunctionality.

A second interesting finding was the selective nature of the ability of IL-7 to partially reverse the suppressive effects of dexamethasone on lymphocytes and restore IFN-γ production. However, IL-7 was not capable of reversing dexamethasone mediated suppression of LPS or resiquimod induced TNF-α production (data not shown). These contrasting effects of IL-7 are likely due to the expression of IL-7 receptor by lymphocytes, whereas monocytes and neutrophils (the cells chiefly responsible for production of TNF-α after TLR stimulation) do not express IL-7 receptor. Thus, the use of dexamethasone or other corticosteroids in patients with sepsis will not necessarily exclude concomitant therapeutic use of IL-7. Theoretically, IL-7 administration to septic patients who are immune suppressed could restore lymphocyte function and not blunt any potential beneficial effect of dexamethasone on the innate immune response. This ability of IL-7 to stimulate lymphocyte function despite concomitant use of corticosteroids is supported by the surprising recent report that dexamethasone potentiates CAR-T cell persistence and function by enhancing IL-7 receptor alpha expression (22). It is important to note, however, that treatment with much higher doses of dexamethasone or comparable corticosteroids might have more potent immunosuppressive effects to blunt the ability of IL-7 to stimulate lymphocytes because of numerous other potent dose-dependent effects of corticosteroids on cell nuclear function (23).

One surprising finding in the current study was the lack of effect of anti-PD-1 mAb to improve IFN-γ production in septic patients. Previously our group reported that anti-PD-1 mAb increased IFN-γ production (24). There are several potential reasons for the disparate findings in the two studies. In the present study, the ELISpot IFN-γ assay was performed on diluted whole blood collected using sodium heparin as the anticoagulant. In the previous study, ELISpot was performed on peripheral blood mononuclear cells (PBMCs) collected using EDTA as the anticoagulant. Although calcium stores were replenished by incubation of the PBMCs in calcium containing media, cytokine production in the PBMCs may have been altered versus immune cells stimulated in diluted whole blood. In this regard, work from our laboratory has shown that collecting patient blood using EDTA as the anticoagulant alters the IFN-γ production compared to heparin as the anticoagulant (unpublished findings). EDTA chelates extracellular and intracellular calcium stores, which are essential for blood coagulation and other cell processes including cell signaling, intracellular trafficking, and transport. Our group previously reported that drugs which alter calcium homeostasis can potently effect LPS-induced cytokine production (25). A second potential reason for the difference in the ability of anti-PD-1 mAb to stimulate IFN-γ production in septic patient blood samples may be due to differences in collection protocol. In the present study, the blood samples were run promptly after collection from the patients. In the previous study, residual blood left over in the hospital hematology laboratory was used. These blood samples were frequently not processed for ELISpot until 24-48 hours after collection. Another possibility may involve immunosuppressive effects of the neutrophils. Several groups have reported that low density neutrophils have potent immunosuppressive effects on lymphocytes (26). Low density neutrophils would be present in the diluted whole blood samples but not present in the PBMC preparation.

The present study adds to the growing consensus that a test of patient immune function is necessary for ideal conduct of clinical trials testing immune adjuvant therapies in sepsis (5). ELISpot may not only enable identification of patients who are severely immune suppressed and candidates for immune adjuvant therapies to restore immune homeostasis, but also allow for titration of the immune drug therapies. A recent study by Traska and colleagues demonstrated that IFN-γ ELISpot was beneficial in guiding immunosuppressive therapy in patients who underwent liver transplantation (27). The investigators quantitated the level of antiviral T-cell activity in PBMCs by measuring IFN-γ secretion upon stimulation with CMV-specific peptide pools. Their results showed that transplant patients who were on triple immune suppressive therapy and had a blunted CMV ELISpot IFN-γ immune response experienced a significantly higher incidence of CMV reactivation. These data suggest prophylactic antiviral therapy effective against CMV should be considered for these patients. Similarly, we believe IFN-γ ELISpot could be useful in identifying septic patients who might benefit from corticosteroid therapy versus those who might be harmed by further immune suppression with corticosteroids.

### Limitations

We speculate that the results of a diluted whole blood ELISpot assay conducted *ex vivo* on septic patient blood sample will be a good indicator of the potential efficacy of immune modulatory drugs administered to patients therapeutically. But we also acknowledge that the ELISpot results may not accurately reflect what happens in patients being treated with the same drugs. It will be important to conduct ELISpot tests in patients who actually receive the immune modulatory drugs in clinical trials. A second limitation is that only a limited number of immune modulatory drugs were tested in the present study. It will be important to evaluate other drugs that hold promise as immune therapies in sepsis. Evaluation of GM-CSF is underway, and results of these studies will be important given the ongoing/planned clinical trials of GM-CSF in pediatric and adult sepsis patients. Another limitation of ELISpot is that the present results reflect overnight stimulation of the patient blood sample to functionally assess the cytokine producing potential of circulating leukocytes. Clinicians may want to begin therapies more quickly instead of waiting for results of an overnight assay. Currently, our team is evaluating shorter incubation times of 4-6 hours and preliminary results are encouraging.

### Conclusion

ELISpot demonstrated excellent dynamic range consistent with the ability of the assay to determine whether the sepsis patient is exhibiting a hyper-inflammatory or immunosuppressive response to the infection at the individual single cell level. Information regarding the relative robustness of the patients’ immune response to sepsis should help guide clinicians in their decision of whether to use drugs to down– or up-regulate patient immunity. Furthermore, the ability of ELISpot to determine the effect of specific immuno-modulatory drugs to independently regulate the innate and adaptive host response could enable precision-based immune drug therapies in sepsis. In addition to sepsis, it is likely that ELISpot may be useful in the management of other diseases (e.g., autoimmunity), where immunomodulatory drugs are widely employed. Although there are a growing number of tests being developed to measure immune cell *function* as a guide to administration of sepsis therapies, we believe ELISpot is particularly attractive given its many capabilities.

---

Supplemental Table 1.

**Supplemental Table 1**
| Stimuli | Final Concentration in Well | Manufacturer (catalog #) |
| --- | --- | --- |
| Anti-CD3/Anti-CD28 | 0.5/5 ug/mL | BioLegend (#300332 and #302934, respectively) |
| Low Anti-CD3/Low Anti-CD28 | 0.125/1.25 ng/mL |  |
| LPS | 1 ng/mL | ENZO Life Sciences (#ALX-581-013-L002) |
| Resiquimod (R848) | 1 ug/mL | InvivoGen (#trl-r848-5) |
| IL-7 | 50 ng/mL | R&D Systems (#207-IL-050) |
| Anti-PD-1 | 1 ug/mL | Bristol Myers Squibb (#NDC 0003-3772-11) |
| Dexamethasone | 1 ug/mL | Mylan (#NDC 67457-423-00) |

**Supplemental Figure 1:**
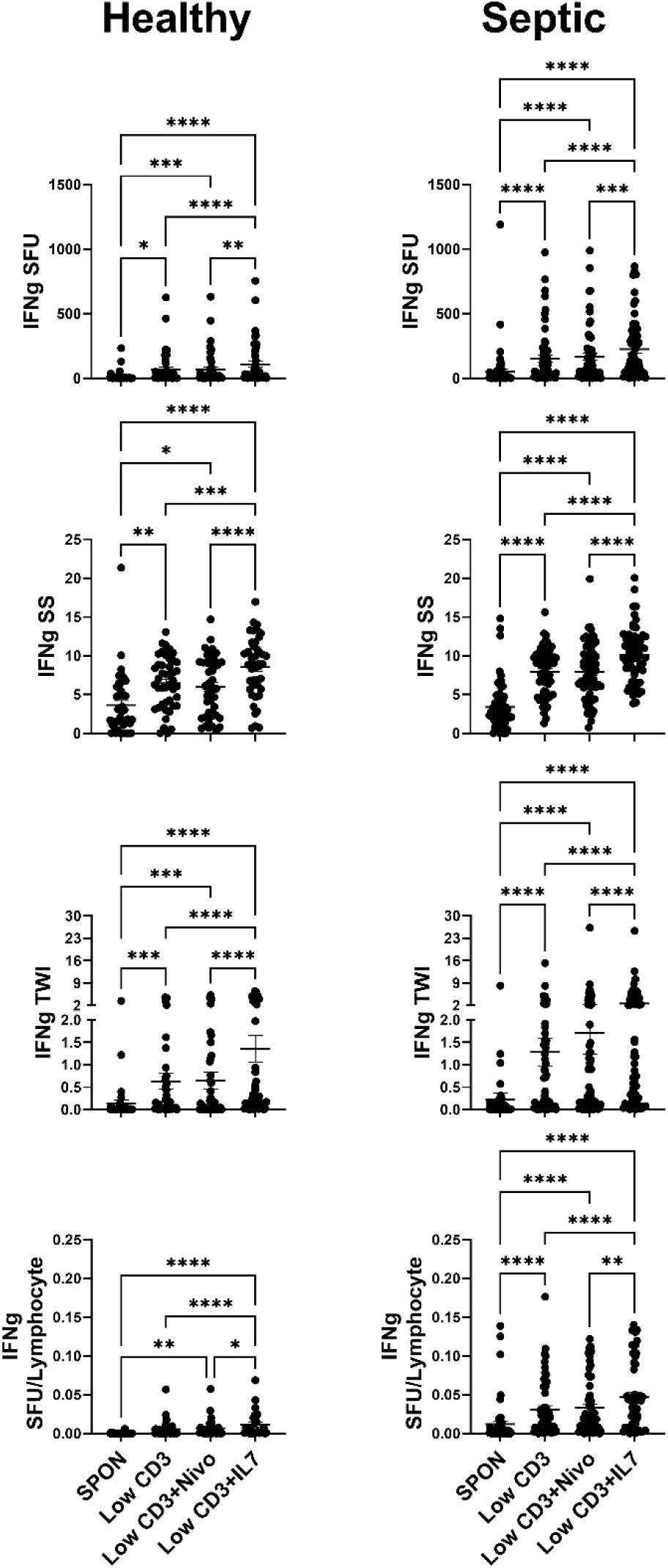
Adjuvant effect on IFNγ production in conjunction with low anti-CD3/CD28 mAb stimulation. Samples were stimulated with low concentration anti-CD3/CD28 mAb with and without either anti-PD-1 mAb (nivolumab; NIVO) or IL-7. Low anti-CD/anti-CD28 (Low CD3) increased IFNγ SFU (upper panels), SS (upper-middle panels) and TWI (lower-middle panels) in samples from both Healthy donors (left) and Septic patients (right) as compared to unstimulated (spontaneous; SPON) production. NIVO did not increase Low CD3 induced IFNγ SFU, SS, or TWI, while IL-7 did increase CD3 induced IFNγ SFU, SS, and TWI. SFU (Top panels) were normalized to the number of lymphocytes plated in the ELISpot well (Bottom panels). Lymphocyte number per well was determined by multiplying the draw specific ALC (K/cumm) by 1000 (converting K/cumm to cells/cumm) then by the volume of blood added to the well (5 µL). Pairwise statistical relationships of SFU/Lymphocyte remain largely the same as SFU. *p<0.05, **p<0.01, ***p<0.001, ****p<0.0001.

**Supplemental Figure 2:**
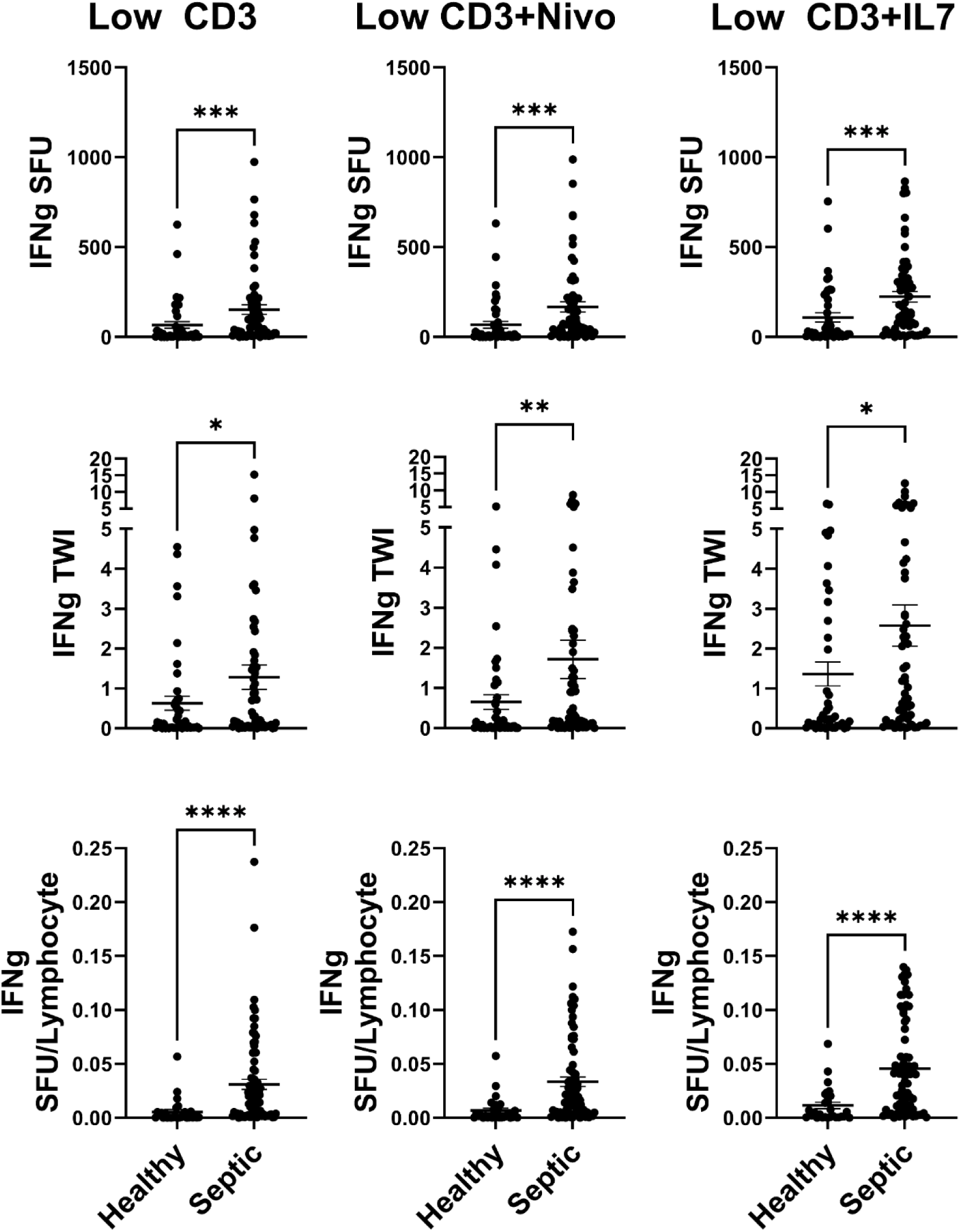
Septic patient samples produce greater IFNγ SFU, TWI, and SFU/Lyphocyte than Healthy donor samples with Low anti-CD3/CD28 mAb stimulation. Comparison between Healthy donor and Septic patient whole blood samples shows septic blood samples have higher IFNγ SFU (top row) than healthy donor samples in response to low anti-CD3/CD28 mAb (CD3) alone (left), low CD3+anti-PD-1 mAb (nivolumab; NIVO; middle), and low CD3+IL-7 (right). To account for high variability in cell counts among septic patients and between septic and healthy samples, SFU were normalized to number of lymphocytes plated in ELISpot wells (bottom row). When SFU were normalized to lymphocyte numbers, the differences between septic patient and healthy donor IFNγ production became even more apparent. As no significant differences were discerned between Low CD3 treated samples and Low CD3+NIVO treated samples, the differences displayed in the “CD3+NIVO” column between healthy and septic are a consequence of the greater responsiveness of septic samples to Low CD3 treatment. *p<0.05, **p<0.01, ***p<0.001, ****p<0.0001.

**Supplemental Figure 3:**
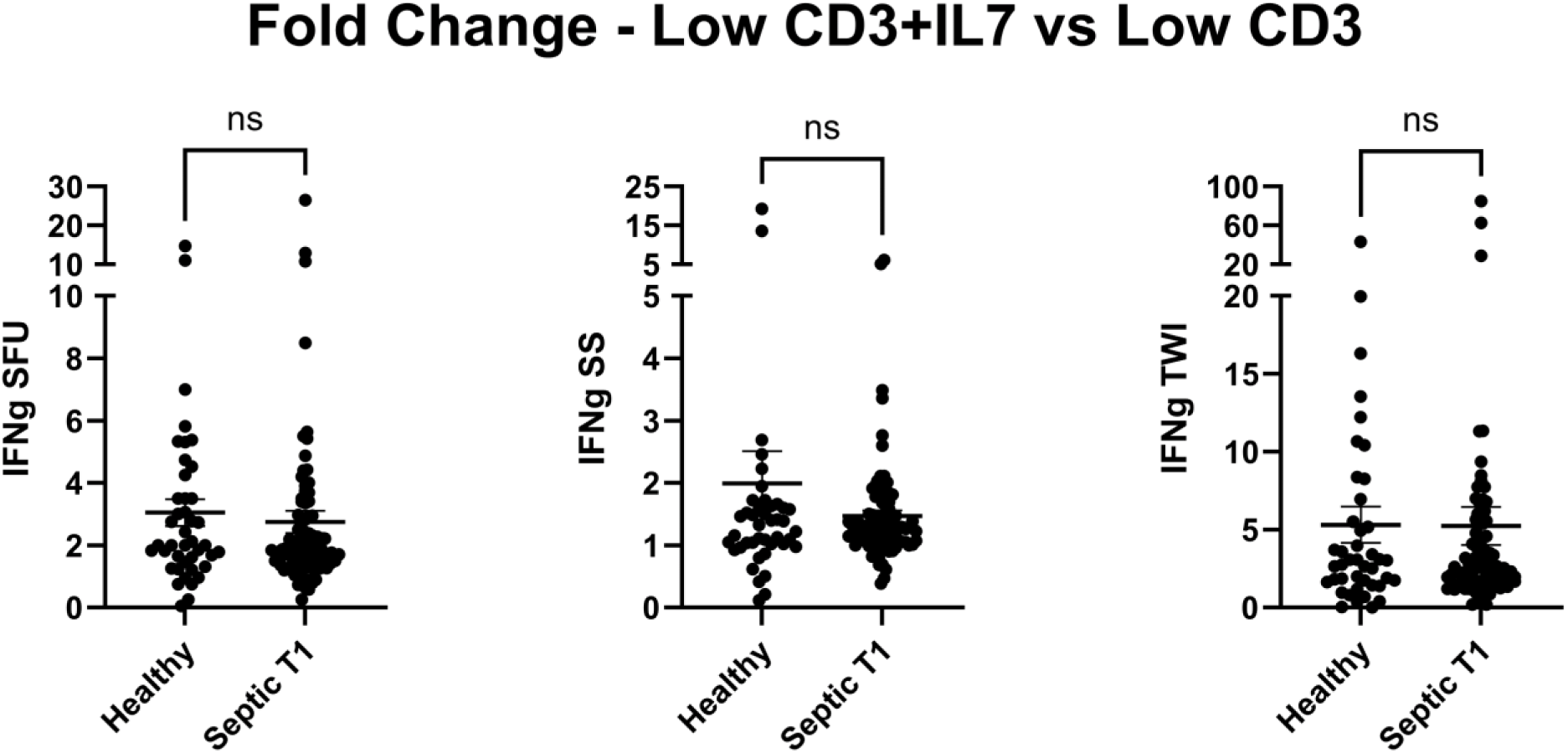
Difference in relative IFNγ response to IL-7 between healthy donors and septic patients when normalized against Low anti-CD3/CD28 mAb stimulation alone. IL-7 fold-changes were calculated for SFU, SS, and TWI for all samples; Low anti-CD3/CD28 mAb (CD3)+IL-7 values were divided by Low CD3 alone to yield fold-change. Unlike with regular CD3 stimulation, there was no difference in responsiveness to IL-7 in the SFU (Left), SS (Middle) or TWI (Right) measurements, based on fold-change, between healthy donors and septic patients. *p<0.05, **p<0.01, ***p<0.001, ****p<0.0001.

**Supplemental Figure 4:**
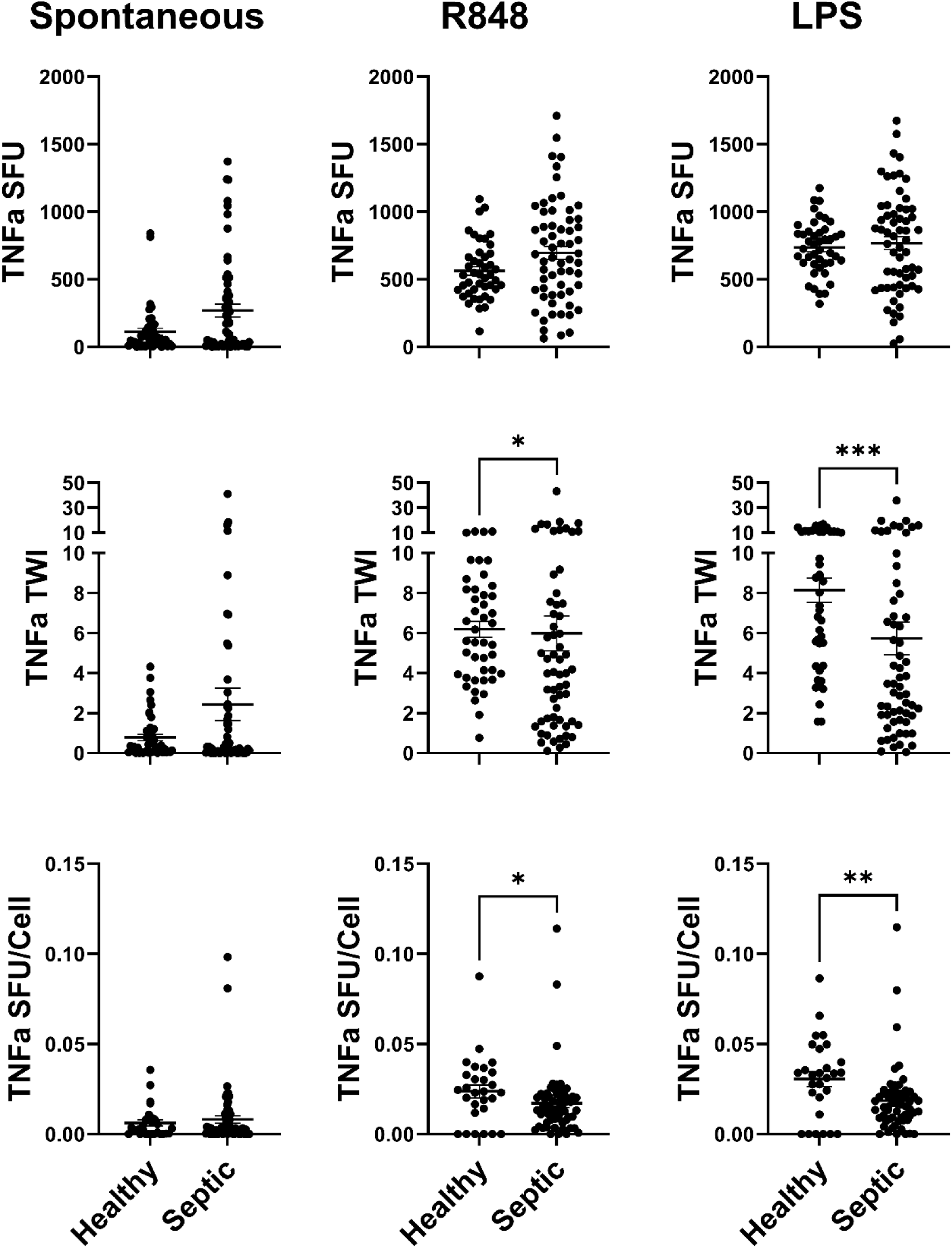
Comparison of TNFα production between healthy and septic samples. Samples from healthy donors and septic patients similarly produced TNFα when unstimulated (spontaneous; SPON; Left Side). They also produced TNFα similarly in response to R848 (Middle) and LPS (Right). *p<0.05, **p<0.01, ****p<0.0001

